# Tunable Elliptical Cylinders for Rotational Mechanical Studies of Single DNA Molecules

**DOI:** 10.1101/2024.09.25.614944

**Authors:** Yifeng Hong, Fan Ye, Xiang Gao, James T. Inman, Michelle D. Wang

## Abstract

The angular optical trap (AOT) is a powerful technique for measuring the DNA topology and rotational mechanics of fundamental biological processes. Realizing the full potential of the AOT requires rapid torsional control of these processes. However, existing AOT quartz cylinders are limited in their ability to meet the high rotation rate requirement while minimizing laser-induced photodamage. In this work, we present a novel trapping particle design to meet this challenge by creating small metamaterial elliptical cylinders with tunable trapping force and torque properties. The optical torque of these cylinders arises from their shape anisotropy, with their optical properties tuned via multilayered SiO_2_ and Si_3_N_4_ deposition. We demonstrate that these cylinders can be rotated at about 3 times the rate of quartz cylinders without slippage while enhancing the torque measurement resolution during DNA torsional elasticity studies. This approach opens new opportunities for previously inaccessible rotational studies of DNA processing.

## INTRODUCTION

Optical tweezers or optical trapping is a cornerstone technique for single-molecule studies of DNA-based processes(*1, 2*). They have been used to establish the elastic properties of a DNA substrate, locate a bound protein on DNA, and track the motions of motor proteins along DNA at high spatial resolution. These techniques have been broadly used to study fundamental biological processes, such as transcription(*3, 4*), replication(*5–8*), recombination(*9*), DNA repair(*10, 11*), and chromatin dynamics(*12–14*). However, a conventional optical trap cannot readily be used to study rotational motions of DNA-based processes, although rotational motions are inherent due to the right-handed helical structure of the DNA (a helical pitch of 10.5 bp or 3.5 nm). For example, as a DNA-based motor protein, such as an RNA polymerase, translocates 10.5 bp, it must also rotate relative to the DNA by one turn(*15*). The next generation of optical trapping, the angular optical trap (AOT), overcomes this limitation by enabling real-time rotation and torque detection(*16, 17*). Since its inception, the AOT has been instrumental in studying the torsional properties of DNA supercoiling(*17–21*) and its impact on fundamental DNA-based processes(*22–25*).

A distinguishing feature of the AOT is its trapping particle. Unlike a conventional optical trap that traps an optically isotropic microsphere, an AOT traps a birefringent cylinder, typically made of quartz, to which a molecule of interest is attached(*17*). The cylinder is nanofabricated to have the extraordinary optical axis perpendicular to the cylindrical axis. When trapped, the cylinder aligns its cylindrical axis with the direction of trapping beam propagation and can be rotated via the rotation of the linear polarization of the incoming beam. The bottom of the cylinder is functionalized for attachment to a biological molecule of interest, such as DNA. Therefore, the cylinder can be used to twist DNA while simultaneously measuring the torque on the DNA(*17*), making the AOT an ideal tool for studying DNA torsional mechanics and DNA-based processes.

The AOT is ideally suited to introduce or relax torque in the DNA by responding to dynamic topological processes on the DNA. For example, during DNA replication, the replisome rotates relative to DNA at about 20-50 turns/s in bacteriophages(*26–28*), 50-100 turns/s in bacteria(*29–31*), and 3-10 turns/s in eukaryotes(*32–34*). Applications of the AOT to study these processes requires the ability to rapidly rotate the cylinder. However, existing quartz cylinders are limited in their rotation rate. When the rotational viscous drag torque on the cylinder exceeds the maximum optical trapping torque, the cylinder rotation will slip relative to the trap polarization rotation(*35*). This limitation cannot be circumvented by simply increasing the laser power since biological samples are prone to laser-induced photodamage(*36–38*). For quartz cylinders, the angular stiffness and the viscous drag torque are tightly coupled. Smaller quartz cylinders can reduce the viscous drag torque, but this size reduction leads to a reduction in the torque and force generation capacities. Alternatively, the angular stiffness can be enhanced via highly birefringent particles(*39–42*), but their potential in torque measurements has not been fully demonstrated at the single-molecule level. Thus, overcoming these challenges requires a new design strategy.

In this work, we have achieved this goal by creating smaller elliptical cylinders with tunable trapping force and torque properties. Instead of using optical birefringence for optical torque generation as with the quartz cylinders, these elliptical cylinders experience an optical torque via their shape anisotropy because the major axis of their elliptical cross-section tends to align with the laser’s linear polarization (Fig. 1a,b)(*43*). These cylinders are made of a metamaterial that affords an effective index of refraction higher than quartz via alternating layers of SiO_2_ and Si_3_N_4_ (Fig. 1c). Unlike quartz cylinders, the optical anisotropy of these cylinders can be continuously tuned via the eccentricity of the cylinder and the Si_3_N_4_ doping fraction.

**Fig. 1.**
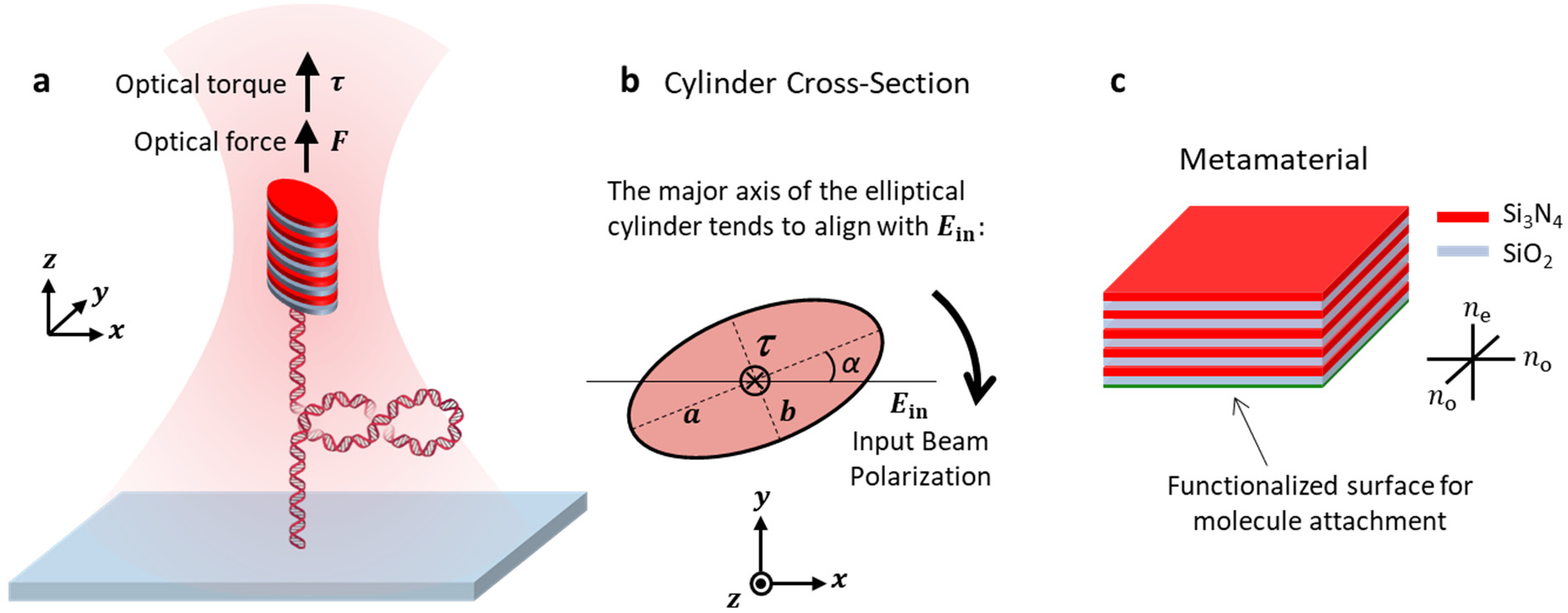
Operational principle of a metamaterial elliptical cylinder in an angular optical trap (AOT). **a.** Experimental configuration of DNA torsional mechanics measurements using a metamaterial elliptical cylinder in an AOT. A DNA molecule, torsionally constrained between the surface of a coverslip of a sample chamber and the bottom of the cylinder, is stretched under a defined force by the optical trap. The input laser beam of the optical trap is linearly polarized. Since the major axis of the cylinder’s elliptical cross-section tends to align with the laser polarization via shape anisotropy, the DNA can be twisted and supercoiled via rotation of the laser polarization. **b.** Optical torque generation of a dielectric elliptical cylinder. When the major axis of the cylinder’s elliptical cross-section is misaligned with input beam polarization forming an angle *β*, a torque is exerted on the cylinder to align the major axis with the polarization. Therefore, the cylinder rotates with the rotation of the polarization. **c.** The metamaterial. The metamaterial is made from periodic stacks of two dielectric materials, SiO_2_ and Si_3_N_4_, with each layer thickness much smaller than the wavelength of the trapping beam and their relative thickness tunable. The resulting material has effective indices of refraction between those of SiO_2_ and Si_3_N_4_ (Fig. S3). Because the material is also birefringent, cylinders made out of such a material also show an extra anti-tilting effect (Fig. S2).

## RESULTS

### Numerical simulations of metamaterial elliptical cylinders

To guide the cylinder design, we performed numerical simulations of trapping force

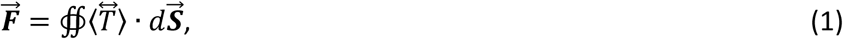

and trapping torque

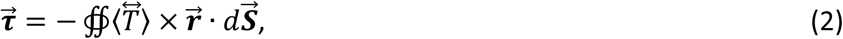

expanding on a numerical platform we previously established(*44*) (Fig. 2a; Materials and Methods), where 〈*T*〉 is the time-averaged Maxwell stress tensor and -*r⃗* is the vector pointing from the center of the cylinder to its surface. Since the viscous drag torque *τ*_drag_ = -*β*_0_θ, where *β*_0_ is the viscous drag coefficient of the cylinder and *θ* is the cylinder’s rotation rate around its cylindrical axis, to reduce the viscous torque, we must reduce *β*_0_, which is roughly proportional to the volume of the cylinder (Fig. S1; Materials and Methods). Our targeting design of cylinder should have a reduced size while affording a trapping force and torque comparable to those of a quartz cylinder which are well suited for DNA torsional mechanical studies and DNA-based processes(*17, 18, 44*).

**Fig. 2.**
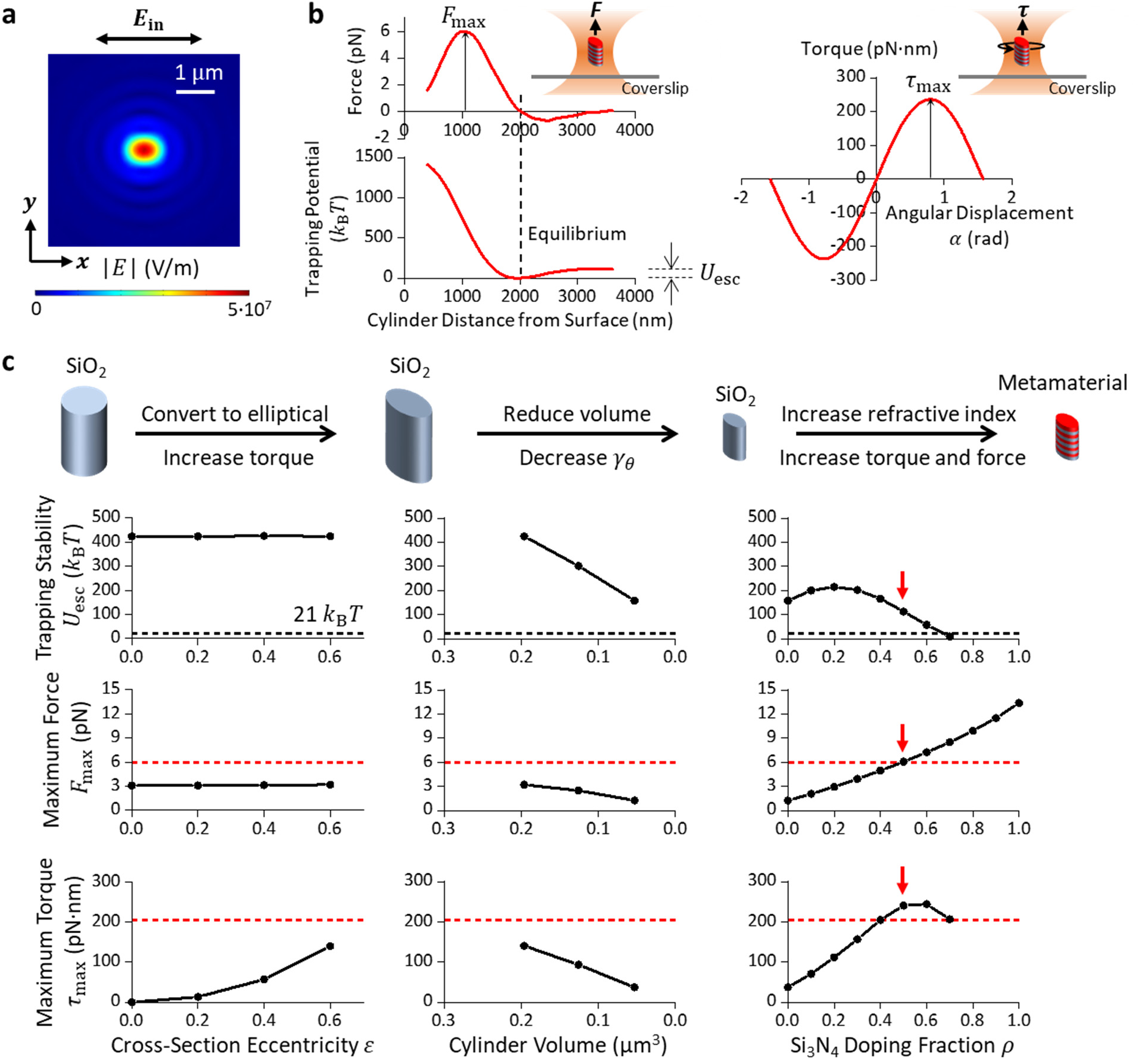
Simulations to guide the design of the metamaterial elliptical cylinder. **a.** Electric field magnitude distribution of a linearly polarized Gaussian beam after being focused to the objective focal plane (NA = 1.3)(*18, 44*). The distribution is elliptical due to the strong focusing effect of a non-paraxial beam. A cylindrical particle with an elliptical cross-section tends to maximize its overlap with the elliptical beam profile for stable angular trapping by aligning its major axis in the direction of light polarization. **b.** Definitions of the maximum trapping force *F*_max_, the trapping stability characterized by the escaping potential *U*_esc_, and the maximum trapping torque *τ*_max_. Shown are example plots of axial force, axial trapping potential, and torque, simulated for a metamaterial elliptical cylinder with major axis length 2*a* = 375 nm, minor axis length 2*b* = 300 nm, height ℎ = 600 nm, and Si_3_N_4_ doping ratio *ρ* = 0.5. **_c._** Simulations characterizing *F*_max_, *U*_esc_, and *τ*_max_ dependencies on cylinder eccentricity, volume, and Si_3_N_4_ doping ratio. To increase the eccentricity of SiO_2_ cylinders, the cylinder height ℎ is kept at 928 nm and the cylinder volume at 0.196 μm^3^. To reduce the volume of SiO_2_ cylinders, the cross-section eccentricity *ε* is kept at 0.6, and the aspect ratio 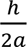 at 1.6. To increase the Si_3_N_4_ doping fraction, the cross-section eccentricity *ε* is kept at 0.6, the cylinder aspect ratio 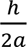 at 1.6, and the cylinder volume at 0.053 μm^3^. The red dashed lines indicate values for the quartz cylinder. The black dashed lines indicate the threshold for the escaping potential. The red arrows indicate values for the targeted metamaterial elliptical cylinders to be fabricated.

We focus on three parameters to guide our design (Fig. 2b): the axial trapping stability characterized by the axial escaping potential barrier height *U*_esc_, the force generation capacity characterized by the maximum force *F*_max_, and the torque generation capacity characterized by the maximum torque *τ*_max_. Note that *U*_esc_ does not need to be exceptionally high as long as it is large enough to ensure stable trapping. Here, we use a threshold of 21 *k*_B_*T* (corresponding to a Boltzmann factor ∼ 10^-9^). For both the simulations and experiments in this work, we have used 30 mW power before the microscope objective, corresponding to 12.6 mW at the specimen plane for an objective with a transmission coefficient of 0.42(*44*). This power has been used in single-molecule measurements without any detectable photodamage to the substrates(*25*).

We first evaluate the torque generation via shape anisotropy by converting the circular cylinders to elliptical cylinders made of an isotropic material. We start the simulation from SiO_2_ (*n* = 1.45 at 1064 nm(*45*)) using circular cylinders with dimensions comparable to those of the quartz cylinders(*17*). As expected, circular cylinders do not generate torque, but conversion to elliptical cylinders allows torque generation. Fig. 2c (left panels) shows how *U*_esc_, *F*_max_, and *τ*_max_ depend on the cylinder cross-section eccentricity 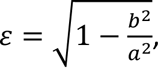, where *a* is the semi-major axis length, and *b* is the semi-minor axis length (Fig. 1b).

Although these elliptical cylinders can generate torque, their viscous drag coefficient *β*_0_is still too large due to their large size (Fig. S1). As shown in Fig. 2c (middle panels), cylinder size reduction (and thus *β*_0_) inevitably reduces the trapping stability, the force generation capacity, and the torque generation capacity, making the cylinders less suitable for single-molecule manipulations and measurements(*18, 25*). To circumvent this issue, we increase the index of refraction of the cylinder material by converting isotropic SiO_2_ to alternating layers of SiO_2_ and Si_3_N_4_ (*n*_Si3N4_ = 2.01(*46*)), which has been shown to increase trapping stiffness(*40, 42*). Each layer thickness is much smaller than the wavelength of the trapping beam. The Si_3_N_4_ doping fraction *ρ* is controlled by varying the relative thickness of the Si_3_N_4_ layers. This multilayered structure behaves as a metamaterial with an effective relative permittivity tensor(*47*):

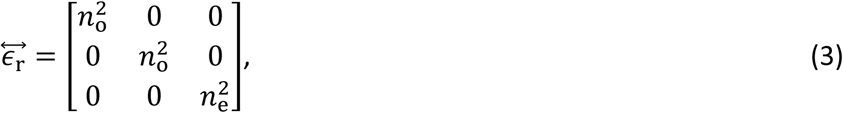

where the index of refraction of the ordinary axis

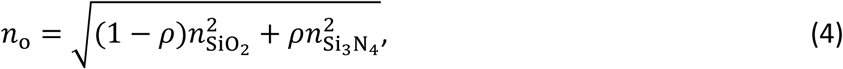

and the index of refraction of the extraordinary axis

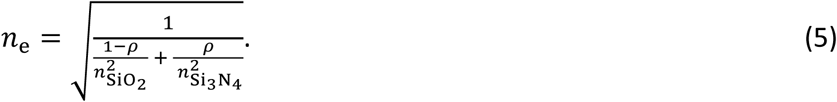

Equations (4) and (5) also show that such a metamaterial is birefringent. Our cylinder design naturally aligns the extraordinary axis along the cylinder axis (Supplementary Materials), which provides more robust angular trapping around the minor axis compared to a cylinder made of an isotropic material of a similar index of refraction(*48*) (Fig. S2). This alignment property of the metamaterial elliptical cylinder is also significantly stronger than that of the quartz cylinder of the same volume tilting around the extraordinary or ordinary axis (Fig. S2).

Thus, the elliptical cylinders made from the metamaterial are more resistant to tilting around this axis, enhancing the angular trapping stability of an AOT.

While increasing the Si_3_N_4_ doping increases force and torque generation capacities, a cylinder with a higher index of refraction also experiences a stronger scattering force, which pushes the cylinder further downstream of the beam waist, consistent with findings from previous studies(*44, 49, 50*). When the doping fraction *ρ* 2’. 0.7, the cylinder can no longer be stably trapped (*U*_esc_ < 21 *k*_B_*T*) (Fig. 2c, right panels). In addition, as the axial equilibrium position of the cylinder relocates further downstream from the beam waist, there is less overlap of the cylinder with the beam profile, reducing in the maximum torque.

### Fabrication of functionalized metamaterial elliptical cylinders

Ultimately, we have chosen the following cylinder parameters for our proof-of-concept fabrication: *ρ* = 0.5 (*n*_e_ = 1.66 and *n*_0_ = 1.75, Fig. S3), 2*a* = 375 nm and 2*b* = 300 nm (eccentricity = 0.6), and cylinder height ℎ = 600 nm. Our simulations indicate that these cylinders should be trapped stably and generate forces and torques comparable to those of the larger quartz cylinder previously used for single-molecule applications(*17, 18, 44*). Importantly, our simulations also indicate that these cylinders have *β*_0_ more than 3 times smaller than that of the quartz cylinders (Fig. S1).

We fabricate the metamaterial elliptical cylinders via a top-down, deep ultraviolet (DUV) lithography process and remove the cylinders with a liftoff method to ensure their uniformity (Fig. 3a; Supplementary Materials). To begin the fabrication, a ∼ 100-nm Al_2_O_3_ film is first deposited as a sacrificial layer(*51*) on a Si wafer (Ultrasil, Lot# 4-14359) via evaporation.

**Fig. 3.**
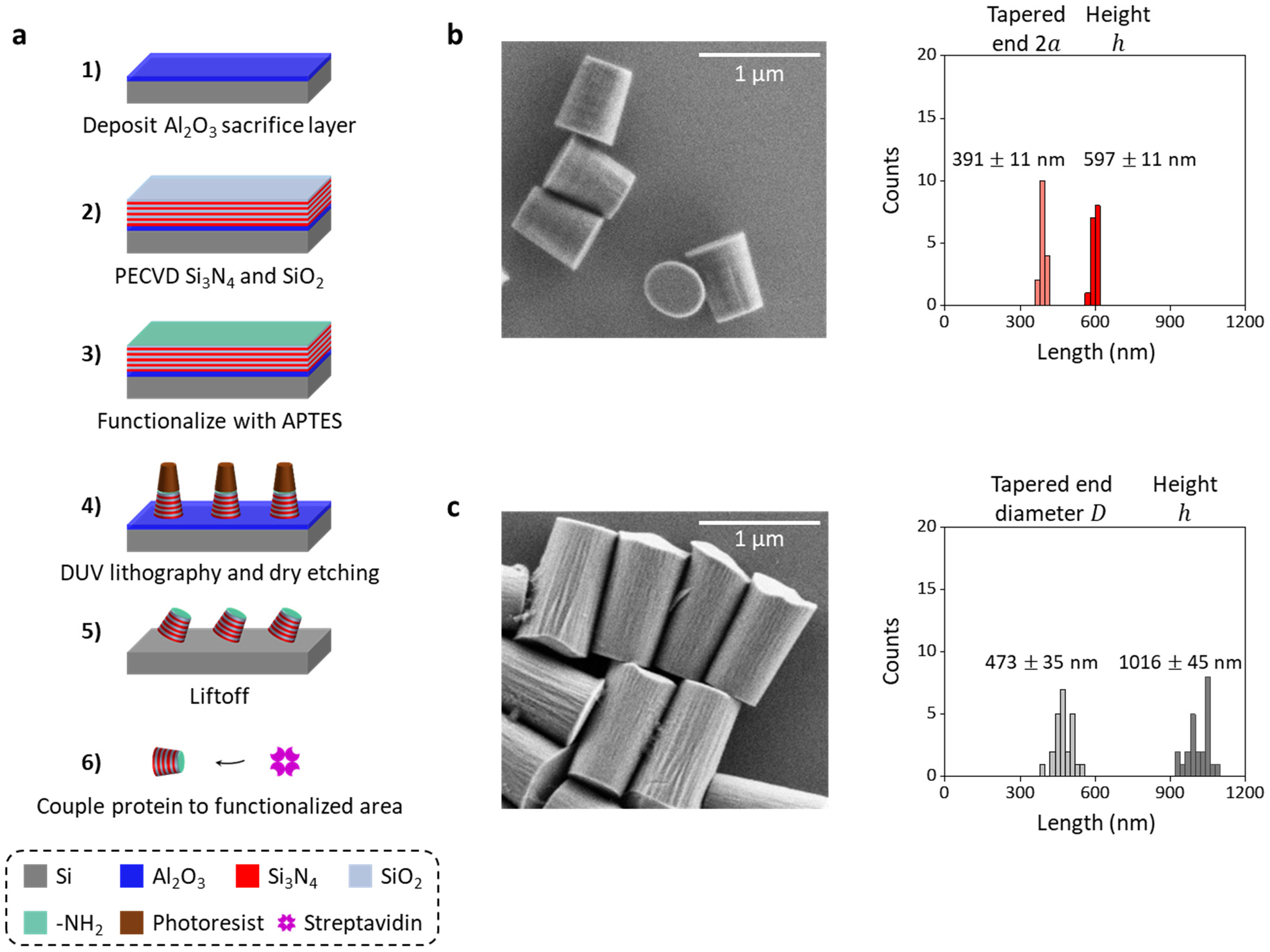
Nanofabrication of metamaterial elliptical cylinders. **a.** Fabrication process flow of metamaterial elliptical cylinders. **b.** An SEM image of nanofabricated metamaterial elliptical cylinders. The size and uniformity characterization is also shown. **c.** An SEM image of nanofabricated quartz cylinders fabricated based on a previously established protocol(*18*). The size and uniformity characterization is also shown.

Subsequently, 5 repeats of alternating layers of Si_3_N_4_ and SiO_2_ are deposited with the plasma-enhanced chemical vapor deposition (PECVD) process, with each layer thickness about 60 nm (<< *λ*_0_ = 1064 nm of our trapping laser). The surface of the sample is then activated with a 20- min O_2_ plasma and then functionalized with (3-aminopropyl)triethoxysilane (Sigma-Aldrich, CAT# 440140) solution(*17*). The elliptical cylinder array is patterned through DUV lithography using a developable antireflection coating (ARC) DS-K101 and photoresist UV-210-0.6. Then, the patterned sample is dry-etched until reaching the Al_2_O_3_ layer. The remaining photoresist and ARC are stripped within heated Microposit Remover 1165 along with sonication. After the photoresist and ARC are removed, the sample is placed in AZ 726 MIF developer for ∼ 3 hr to liftoff the cylinders. Finally, the cylinders are collected through a spin-down using a centrifuge and then coupled to streptavidin (Agilent, CAT# SA-10) with glutaraldehyde (Sigma-Aldrich, CAT# G5882) as the linker(*17*).

The resulting metamaterial elliptical cylinders have the following dimensions based on scanning electron microscope (SEM) images (Fig. 3b): tapered-end 2*a* = 391 ± 11 nm and ℎ = 597 ± 11 nm (mean ± s.d., *N* = 16), compared to those of the fabricated quartz cylinders: tapered-end diameter 473 ± 35 nm and ℎ = 1016 ± 45 nm (mean ± s.d., *N* = 24) (Fig. 3c)(*44*). The metamaterial elliptical cylinders represent about a 3-fold volume reduction compared with the quartz cylinders. In addition, the use of the lift-off method greatly reduces the height variations by about 4-fold.

### Enhanced maximum rotation rate via metamaterial elliptical cylinders

When we placed these cylinders in the AOT, we found that they can be stably trapped, as predicted by the simulations. We show that our metamaterial elliptical cylinders can generate a maximum force and a maximum torque comparable to the quartz cylinders (Fig. 4a,b), consistent with our prediction. In addition, we measured the rotational motion of metamaterial elliptical cylinders and found they show a 3-fold reduction in *β*_0_: 3.2 ± 0.3 pN·nm·s/turn (mean ± s.d., *N* = 17), in comparison to 9.4 ± 1.7 pN·nm·s/turn (mean ± s.d., *N* = 14) for the quartz cylinder (Fig. 4a). The reduced *β*_0_should result in a faster cylinder rotation rate without slippage with the metamaterial elliptical cylinders.

**Fig. 4.**
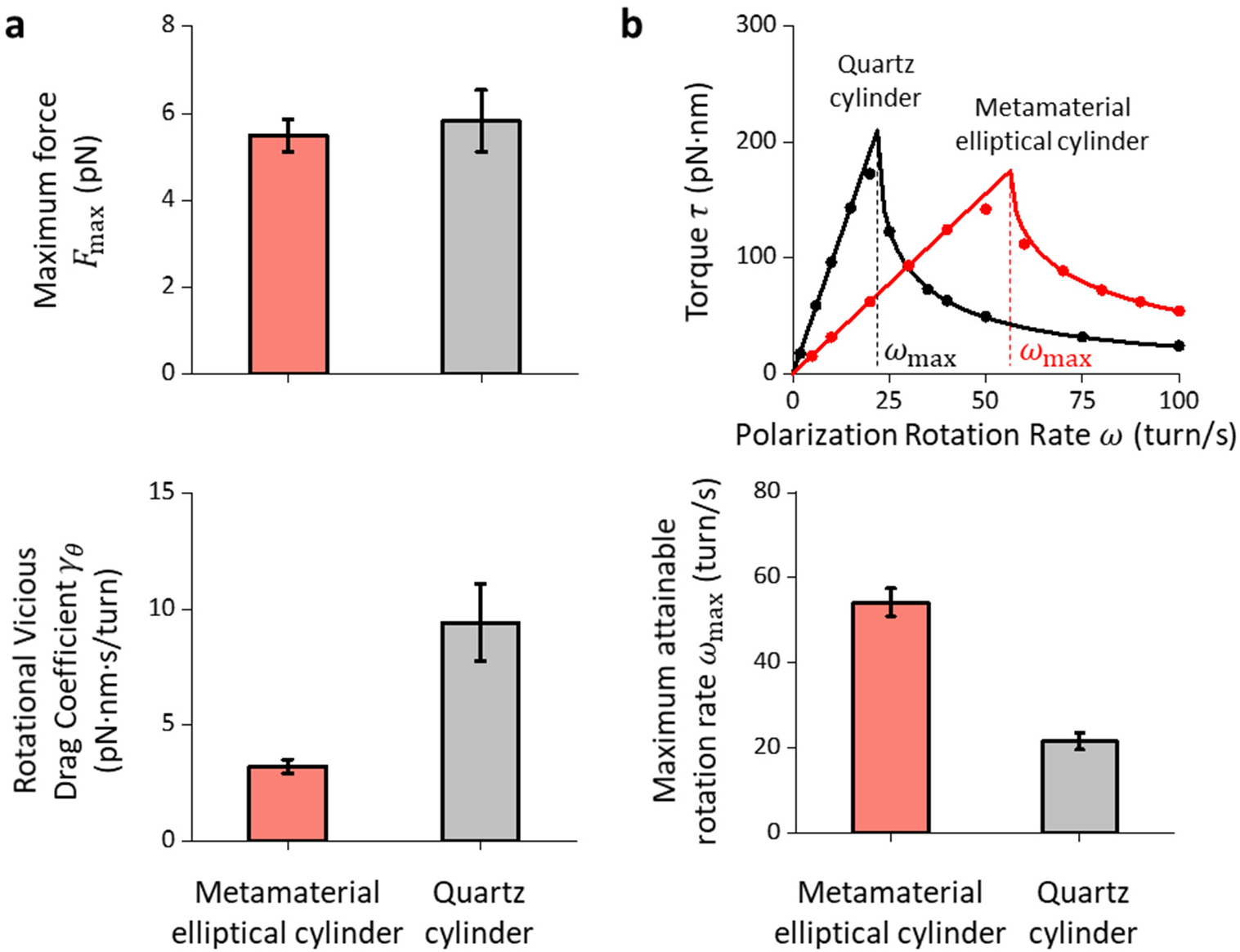
Trapping properties and maximum rotation rate of metamaterial elliptical cylinders. **a.** Measurements of the maximum trapping force *F*_max_(top panel, *N* = 14 metamaterial elliptical cylinder; *N* = 14 quartz cylinders) at 30 mW laser power before the objective and rotational viscous drag coefficient *β*_0_(bottom panel, *N* = 17 metamaterial elliptical cylinders; *N* = 14 quartz cylinders). Values shown are mean ± s.d. **b.** Method to determine the maximum trapping torque *τ*_max_ and the maximum rotation rate ω_max_ without slippage at 30 mW laser power before the objective. Top panel: We measure the mean viscous torque on the cylinder *τ* (which also indicates the cylinder rotational rate against the rotational viscous drag) as a function of the polarization rotation rate *ω* (dots). The torque can reach a maximum *τ*_max_ at the maximum rotation rate *ω*_max_ without slippage. W solid lines) to *τ* = *β*_0_*ω* for *ω* ≤ *ω*_max_ and *τ* = 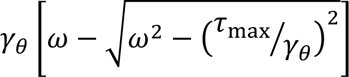 for *ω* > *ω* to determine *τ*_*max*_ and *ω*_max_ (*35*). Bottom panel: Measurements of the maximum rotational rate without slippage *ω*_max_ (*N* = 9 metamaterial elliptical cylinder; *N* = 8 quartz cylinders). Values shown are mean ± s.d.

To directly determine the maximum rotation rate without slippage *ω*_max_, which is expected to be related to *τ*_max_ and *β*_0_:

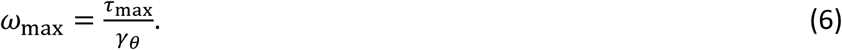

We measured the torque due to the viscous rotational drag as a function of the polarization rotation rate *ω* using previously established methods(*35*). As shown in Fig. 4b (top panel), when ≤ *ω*_max_, the cylinder follows the polarization rotation rate, and the viscous drag torque increases linearly. When *ω* > *ω*_max_, the cylinder no longer tracks the polarization and rotates slower than the polarization rotation rate, leading to a decrease in the viscous drag torque with an increase in the polarization rotation rate.

We found that *ω*_max_ for the metamaterial elliptical cylinders is about 3 times that of the quartz cylinders at a given laser power (Fig. 4b, bottom panel). Alternatively, if the cylinders are rotated at the same rate, the trapping power can be reduced by about 3 times for the metamaterial elliptical cylinders without slippage. Previous studies showed that the near-infrared trapping laser light can be significantly absorbed by cellular components(*36, 37*) and proteins and DNA(*38*), inducing photodamage to biological molecules. Thus, power reduction using metamaterial elliptical cylinders enables the opportunity for monitoring fast DNA rotation, for instance, by the bacteria replisome (50-100 turn/s(*29–31*)), with reduced irreversible photodamage to the sample.

For comparison, we also fabricated isotropic SiO_2_ elliptical cylinders (eccentricity = 0.6) that are of similar size as the quartz cylinders and calibrated them. Compared with the quartz cylinders, they have about 60% of the maximum force, 50% of the maximum torque, and a similar *β*_0_ (Fig. S4), demonstrating the need to increase the index of the refraction and reduce the size to increase the maximum rotation rate without slippage at a given power.

### Enhanced single-molecule torque resolution via metamaterial elliptical cylinders

In addition, the reduced *β*_0_ of the metamaterial elliptical cylinders also have an extra benefit of a greater signal-to-noise ratio (SNR) in the torque measurement of a DNA molecule:

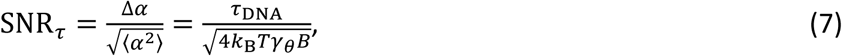

where *α* is the angular displacement of the cylinder from its angular equilibrium, *τ*_DNA_ is the DNA torque to be measured, *k*_B_*T* is the thermal energy, and *B* is the measurement bandwidth (which is assumed to be significantly smaller than the particle’s corner frequency)(*1*). Based on eq. (7), we estimate that when the DNA generates about 10 pN·nm torque, which can stall a transcribing RNA polymerase(*22*), the SNR of measurements with a quartz cylinder is around 2. The 3-fold reduction in *β*_0_for the metamaterial elliptical cylinders should provide a 1.7-fold reduction in the noise of the measured torque of a DNA molecule.

To assess this, we held a DNA molecule under a constant force using a metamaterial elliptical cylinder, twisted the DNA with the AOT, and then directly measured the torque required to twist DNA (Fig. 5; Materials and Methods). To enable accurate force detection and exertion, we corrected the small but non-negligible trap-height dependent force offset due to the Fabry-Perot effect due to backscattered trapping light (Fig. S5). We found that metamaterial elliptical cylinders can accurately measure the torsional properties of DNA under different forces (Fig. S6).

**Fig. 5.**
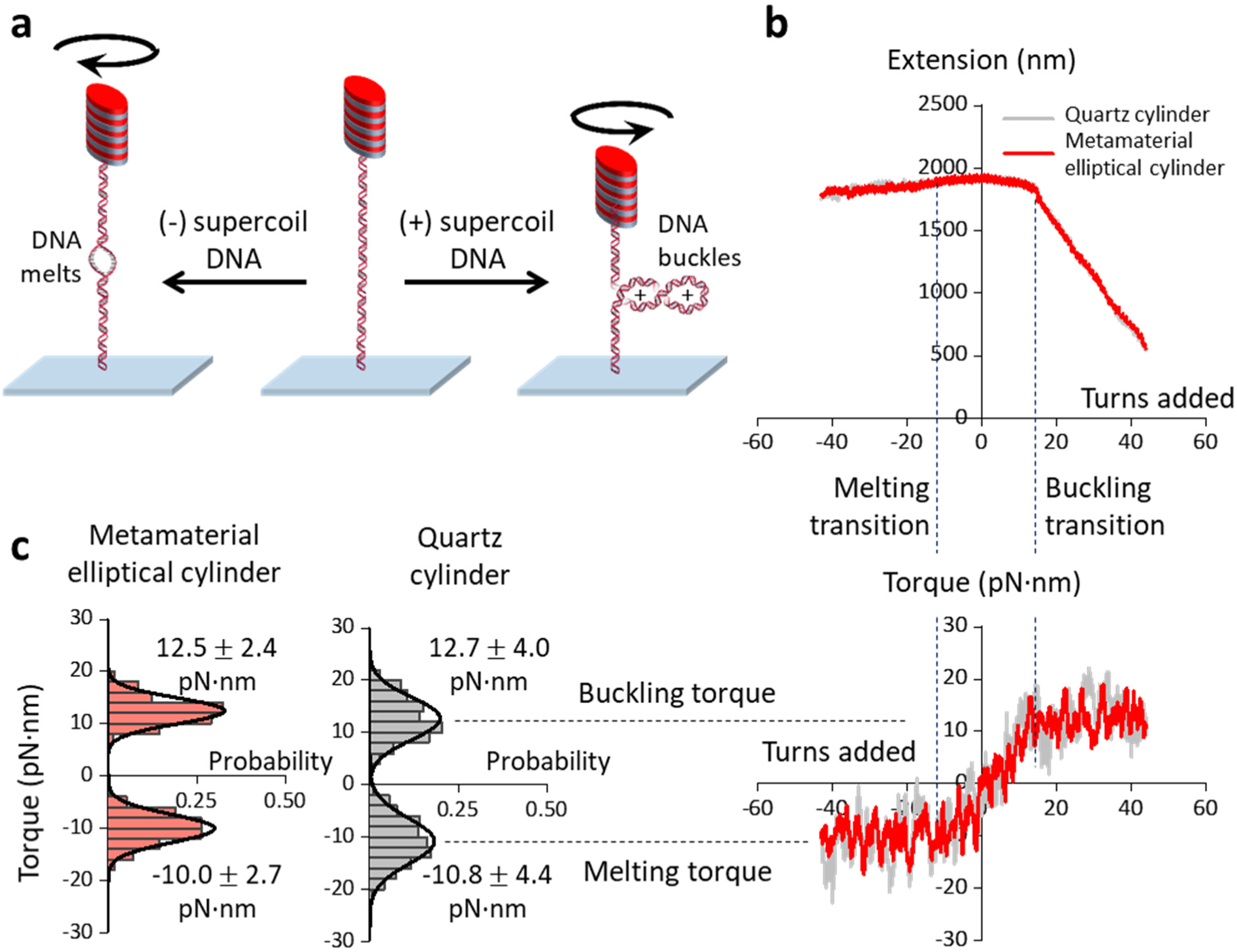
DNA torque measurements using a metamaterial elliptical cylinder, in comparison with those from a quartz cylinder. **a.** Experimental configuration for the measurements. A 6.5 kb DNA molecule was torsionally constrained between the cylinder and the surface and was held under a constant force. The DNA molecule was either positively supercoiled to buckle and extrude a plectoneme or negatively supercoiled to induce melting. **b.** Measured DNA extension and torque as a function of turns added to DNA under 1 pN force. The cylinder was rotated at 2 turn/s in either direction. The data were collected at a sampling rate of 400 Hz by the AOT and smoothed to 0.1 turn for the extension data and 1.0 turn for the torque data. **c.** Histograms of measured torque upon (+) DNA buckling and (-) DNA melting. Each histogram is fit by a Gaussian, with the mean and the s.d. of the fit shown.

Figure 5b shows the DNA extension and torque measurements from a single molecule of DNA held under 1 pN force. As expected(*18–21, 25*), when DNA is positively supercoiled, the DNA extension remains nearly constant and the torque increases linearly until the DNA buckles. After the DNA is buckled, further (+) DNA supercoiling results in a linear decrease of the DNA extension and extrusion of a plectoneme while the buckling torque remains constant. When DNA is negatively supercoiled, the DNA extension remains nearly constant. Further (-) DNA supercoiling leads to a DNA melting transition, beyond which the melting torque remains constant.

We examined the measured torque precisions at the buckled DNA torque plateau (around +12.8 pN·nm) and the DNA melting torque plateau (around -10.5 pN·nm). The torque plateaus measured using a metamaterial elliptical cylinder and a quartz cylinder agree with each other and with previous results(*18*). However, the standard deviations of the torque values of the metamaterial elliptical cylinder are about 1.7-fold smaller than those of the quartz cylinder (Fig. 5c), which is consistent with our prediction. Thus, we demonstrated that these metamaterial elliptical cylinders can facilitate high-resolution single-molecule torque measurements.

## DISCUSSION

Taken together, we demonstrated, both theoretically and experimentally, that our small- size metamaterial elliptical cylinders can permit cylinder rotation about 3 times the rate of the quartz cylinders while providing high force and torque for DNA torsional mechanics studies with enhanced torque resolution. Moreover, our methodology offers versatility in tuning the refractive index, shape anisotropy, and cylinder size to optimize the trapping properties. We anticipate new opportunities to use these cylinders to enable previously inaccessible rotational studies of DNA-based biological processes.

## MATERIALS AND METHODS

### Simulations of Optical Force and Torque

We performed numerical simulations of the optical force and torque on the trapped cylinder to facilitate our design. The simulations were carried out with the wave optics module in COMSOL Multiphysics 5.5. The simulation method was previously developed(*44*), where the tightly focused Gaussian beam was numerically generated with NA = 1.3 and an aperture filling ratio of 0.98 based on our setup(*18*). The spherical aberration induced by the oil-aqueous surface was considered in our simulations. The force simulation was performed by translocating the cylinder across the trap with the cylinder oriented such that there was zero torque. The zero net torque condition was achieved by aligning the beam polarization with a metamaterial elliptical cylinder’s major axis or a quartz cylinder’s extraordinary axis. The torque simulation was performed by rotating the cylinder along with its permittivity tensor around its cylindrical axis. The force and torque were calculated by surface integration of the Maxwell stress tensor.

### Simulations of Rotational Viscous Drag Coefficient

We performed simulations to estimate the rotational viscous drag coefficient *β*_0_ of cylindrical particles (Fig. S1). The simulations were carried out with the laminar flow module in COMSOL Multiphysics 6.1. In our simulation configuration, the cylinder, surrounded with the aqueous solution (density = 1000 kg/m^3^ and viscosity = 0.001 Pa·s)(*52*), was placed in the middle of a cubic box (length of side = 25 µm). The wall of the cylinder was set to have an angular sliding velocity of 1 turn/s. The *β*_0_ was then evaluated using the calculated torque on the cylinder via surface integration of the stress.

### DNA Template Preparation

A 6,546 bp single-labeled DNA was used for the DNA stretching experiment, which was prepared via PCR from plasmid pMDW133 (sequence available upon request) as described previously(*44*). A 6,481 bp multi-labeled DNA was used for the DNA twisting experiment, which was prepared based on a previously published protocol(*44*).

### Data Acquisition and Analysis

All the AOT experiments were performed at room temperature (23 °C). Each sample chamber used for the AOT measurements was assembled from a nitrocellulose-coated coverslip and a Windex-cleaned glass slide(*25*). The DNA tethering was achieved by following a previously reported assay(*18*). All the experiments were performed in phosphate buffered saline (PBS, Invitrogen AM9625: 137 mM NaCl, 2.7 mM KCl, 8 mM Na_2_HPO_4_, 2 mM KH_2_PO_4_, pH 7.4) with 0.6 mg/mL β-casein (Sigma, C6905).

The untethered cylinder spinning experiments were carried out at a laser power of 30 mW (before the objective). Data were acquired and recorded at 10 kHz.

The DNA stretching experiments (Fig. S5) were carried out at a laser power of 30 mW (before the objective). The single-labeled DNA molecule was stretched at a constant velocity of 200 nm/s. Data were acquired at 10 kHz and recorded after averaging to 400 Hz.

The DNA twisting experiments (Fig. 5; Fig. S6) were carried out under a constant-force mode at a laser power of 30 mW (before the objective). The cylinder was rotated at 2 turn/s to supercoil the attached DNA. Data were acquired at 10 kHz and recorded after averaging to 400 Hz. The measurements using quartz cylinders were performed under the same condition.

## Supplementary Materials This PDF file includes

Supplementary Text Figs. S1 to S6

## Acknowledgements

We thank members of the Wang Laboratory for helpful discussions and technical support. We thank Dr. X. Jia and T.M. Kay for the DNA template preparations used in this work. This work was performed in part at the Cornell NanoScale Facility, a member of the National Nanotechnology Coordinated Infrastructure (NNCI), which is supported by the National Science Foundation (Grant NNCI-2025233).

## Author contributions

Y.H., F.Y., and M.D.W conceived the design and methodology. Y.H. and F.Y. performed simulations. Y.H. fabricated the cylinders. Y.H., X.G., and J.T.I. upgraded the AOT software for real-time force-offset corrections. Y.H. performed measurements and data analysis. M.D.W. supervised the project.

## Data and material availability

All data needed to evaluate the conclusions in the paper are present in the paper and/or the Supplementary Materials.

