## Supplementary Materials for "Tunable Elliptical Cylinders for Rotational Mechanical Studies of Single DNA Molecules"

### Cylinders Tilting in the Trapping Beam

The AOT measures the torque on the cylinder about its cylindrical axis, so rotation about the other two orthogonal axes is undesirable. The elongated cylindrical shape is chosen to ensure that a trapped cylinder aligns its cylindrical axis with the direction of the trapping beam propagation. To limit cylinder rotation around the other two axes, the cylinder must have a high angular trapping stiffness around those axes to resist tilting(*1*).

To evaluate the angular trapping stiffness around those two axes, we performed COMSOL simulations using the relative permittivity tensors of the metamaterial elliptical cylinder:

$$\overleftrightarrow{\epsilon}_{r,\theta_x} = \begin{bmatrix} n_\mathrm{o}^2 & 0 & 0 \\ 0 & n_\mathrm{e}^2 \sin^2\theta_x + n_\mathrm{o}^2 \cos^2\theta_x & (-n_\mathrm{e}^2 + n_\mathrm{o}^2)\sin\theta_x\cos\theta_x \\ 0 & (-n_\mathrm{e}^2 + n_\mathrm{o}^2)\sin\theta_x\cos\theta_x & n_\mathrm{e}^2\cos^2\theta_x + n_\mathrm{o}^2\sin^2\theta_x \end{bmatrix}, \tag{S1}$$

$$\overleftrightarrow{\epsilon}_{r,\theta_y} = \begin{bmatrix} n_e^2\sin^2\theta_y + n_o^2\cos^2\theta_y & 0 & (n_e^2 - n_o^2)\sin\theta_y\cos\theta_y \\ 0 & n_o^2 & 0 \\ (n_e^2 - n_o^2)\sin\theta_y\cos\theta_y & 0 & n_e^2\cos^2\theta_y + n_o^2\sin^2\theta_y \end{bmatrix}, \quad \text{(S2)}$$

where $\theta_x$ and $\theta_y$ are the tilting angles around the elliptical cylinder's major axis and minor axis, respectively. Compared to an isotropic elliptical cylinder, a metamaterial elliptical cylinder can enhance the angular stiffness about its minor axis by 3-fold (Fig. S2), resulting in more robust anti-tilting around this axis.

We performed similar simulations for a quartz cylinder, which has the following relative permittivity tensors:

$$\overleftrightarrow{\epsilon}_{r,\theta_x} = \begin{bmatrix} n_e^2 & 0 & 0 \\ 0 & n_o^2 & 0 \\ 0 & 0 & n_o^2 \end{bmatrix}, \quad \text{(S3)}$$

$$\overleftrightarrow{\epsilon}_{r,\theta_y} = \begin{bmatrix} n_e^2\cos^2\theta_y + n_o^2\sin^2\theta_y & 0 & (-n_e^2 + n_o^2)\sin\theta_y\cos\theta_y \\ 0 & n_o^2 & 0 \\ (-n_e^2 + n_o^2)\sin\theta_y\cos\theta_y & 0 & n_e^2\sin^2\theta_y + n_o^2\cos^2\theta_y \end{bmatrix}, \quad \text{(S4)}$$

where the permittivity tensor is independent of the tilting angle $\theta_x$ around the cylinder's extraordinary axis, and $\theta_y$ is the tilting angle around the ordinary axis. When having the same volume, the metamaterial elliptical cylinder is about 2~4-fold more resistant to tilting than the quartz cylinder around its extraordinary axis and ordinary axis (Fig. S2).

**Detailed Fabrication Protocol of Metamaterial Elliptical Cylinders**

1. Clean the Si wafer (Ultrasil, Lot# 4-14359) with a hot piranha solution.
2. Deposit ~ 100 nm $Al_2O_3$ sacrificial layer onto the Si wafer via evaporation.
3. Deposit $Si_3N_4$ then $SiO_2$ with a single layer thickness of ~ 60 nm through plasma-enhanced chemical vapor deposition (PECVD).
4. Repeat step 3 for 5 times, resulting in a total thickness of ~ 600 nm metamaterial.
5. Activate the surface with 20 min $O_2$ plasma.
6. React with (3-Aminopropyl)triethoxysilane solution(*1*).
7. Coat the anti-reflection coating DS-K101, then soft bake at 185 °C for 90 s.

8. Coat the photoresist UV-210-0.6, then soft bake at 135 °C for 90 s.
9. DUV lithography with a photomask of ellipses (eccentricity = 0.6), followed by a post-exposure bake at 135 °C for 90 s.
10. Develop in AZ 726 MIF developer for 60 s.
11. Hard bake at 115 °C for 60 s.
12. $O_2$ descum for 30 s.
13. Dry etch (reactive ion etching) with the chemistry of 45 sccm $CHF_3$, 15 sccm Ar, 50 mTorr, 200 W until reaching the $Al_2O_3$ layer.
14. Remove the photoresist with heated Microposit Remover 1165 along with sonication.
15. Liftoff the metamaterial elliptical cylinders with AZ 726 MIF developer for ~ 3 hr.
16. Collect the cylinder through a centrifuge.

**Detailed Fabrication Protocol of Isotropic $SiO_2$ Elliptical Cylinders**

We also fabricated isotropic $SiO_2$ elliptical cylinders to verify the methodology of using shape anisotropy for torque generation on the AOT. The fabrication protocol of those cylinders is listed as follows:

1. Clean the Si wafer (Ultrasil, Lot# 4-14359) with a hot piranha solution.
2. Deposit ~ 100 nm $Al_2O_3$ sacrificial layer onto the Si wafer via evaporation.
3. Deposit ~ 1 μm $SiO_2$ through PECVD.
4. Activate the surface with 20 min $O_2$ plasma.
5. React with (3-Aminopropyl)triethoxysilane solution(*1*).
6. Coat the anti-reflection coating DS-K101, then soft bake at 185 °C for 90 s.
7. Coat the photoresist UV1400-1.4, then soft bake at 135 °C for 90 s.
8. DUV lithography with a photomask of ellipses (eccentricity = 0.6), followed by a post-exposure bake at 115 °C for 90 s.
9. Develop in AZ 726 MIF developer for 60 s.
10. Hard bake at 110 °C for 60 s.
11. $O_2$ descum for 70 s.

12. Dry etch (reactive ion etching) with the chemistry of 45 sccm $CHF_3$, 15 sccm Ar, 50 mTorr, 200 W until reaching the $Al_2O_3$ layer.
13. Remove the photoresist with heated Microposit Remover 1165 along with sonication.
14. Liftoff the metamaterial elliptical cylinders with AZ 726 MIF developer for ~ 3 hr.
15. Collect the cylinder through a centrifuge.

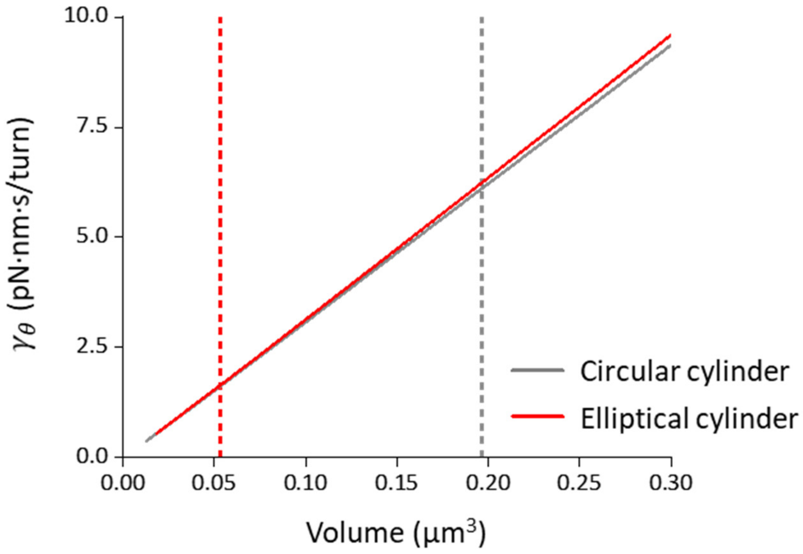


**Fig. S1.** Rotational viscous drag coefficients $\gamma_\theta$ of cylinders.

Simulated viscous drag coefficient $\gamma_\theta$ dependence on the cylinder volume with a fixed shape. For elliptical cylinders, the cross-section eccentricity $\varepsilon$ is kept at 0.6, the cylinder aspect ratio $\frac{h}{2a}$ at 1.6. For circular cylinders, the cylinder aspect ratio $\frac{h}{D}$ is kept at 2. The red dashed line indicates the targeted volume of metamaterial elliptical cylinders to be fabricated. The grey dashed line indicates the typical size of quartz cylinders used for single-molecule studies(*1, 2*).

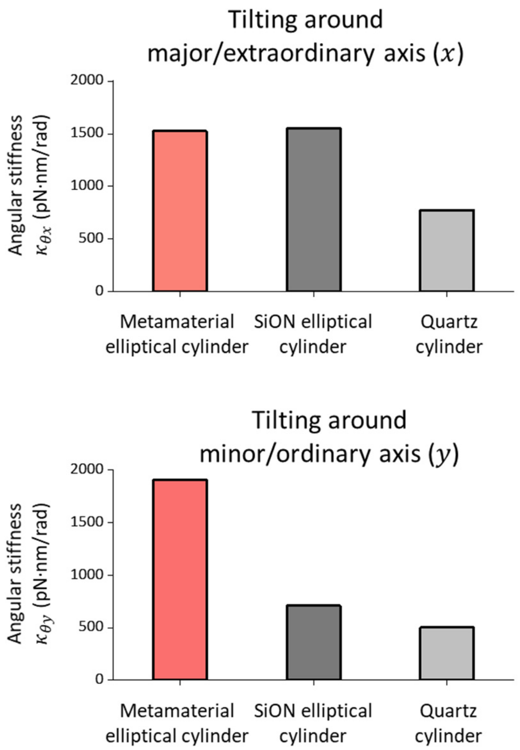


**Fig. S2.** Cylinder tilting in the trapping beam.

Simulated angular stiffness of metamaterial and SiON elliptical cylinders tilting around the major axis ($x$) and the minor axis ($y$). Also shown is simulated angular stiffness of a quartz cylinder tilting around its extraordinary axis ($x$) and ordinary axis ($y$). The volumes of these three types of cylinders are kept the same. The simulated angular stiffness characterizes how robustly cylinders align their cylindrical axis with the direction of the trapping beam propagation. The metamaterial ($n_e$ = 1.66 and $n_o$ = 1.75) and isotropic SiON ($n$ = 1.75) elliptical cylinders both have the dimensions of $\varepsilon$ = 0.6, $\frac{h}{2a}$ = 1.6, and $h$ = 600 nm. The quartz cylinder ($n_e$ = 1.54 and $n_o$ = 1.53)(*3*) has the dimensions of $\frac{h}{D}$ = 1.8 and $h$ = 600 nm.

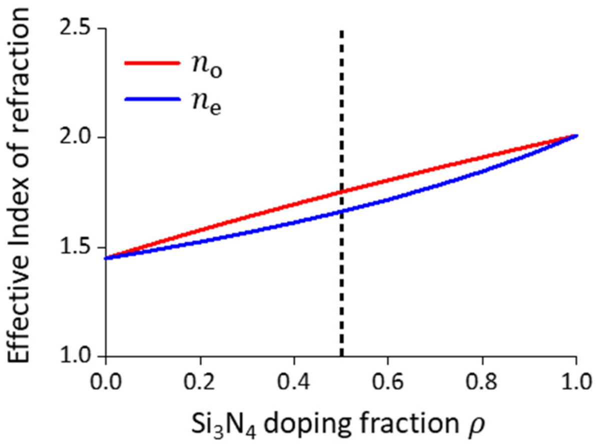


**Fig. S3.** The effective refractive index of the metamaterial doped with $SiO_2$ and $Si_3N_4$. The dashed line indicates the targeted doping fraction of the fabricated elliptical cylinders.

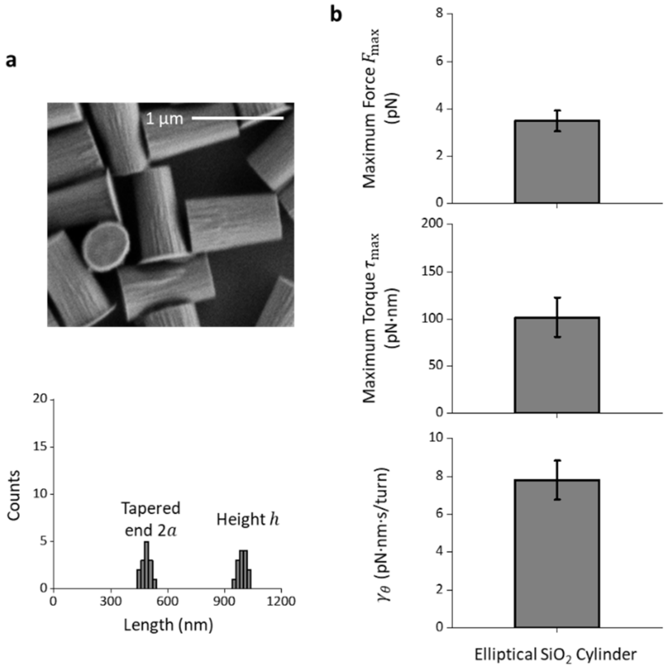


**Fig. S4.** Trapping properties of $SiO_2$ elliptical cylinders**.**

**a**. A scanning electron microscope image of nanofabricated $SiO_2$ elliptical cylinders. Scale bar: 1 µm. The cylinder dimension distributions are also shown.

**b**. Maximum trapping force $F_{\max}$, maximum trapping torque $\tau_{\max}$ at 30 mW laser power before the objective, and rotational viscous drag coefficient $\gamma_\theta$ of the $SiO_2$ elliptical cylinders (*N* = 14 cylinders). Values shown are mean $\pm$ s.d.

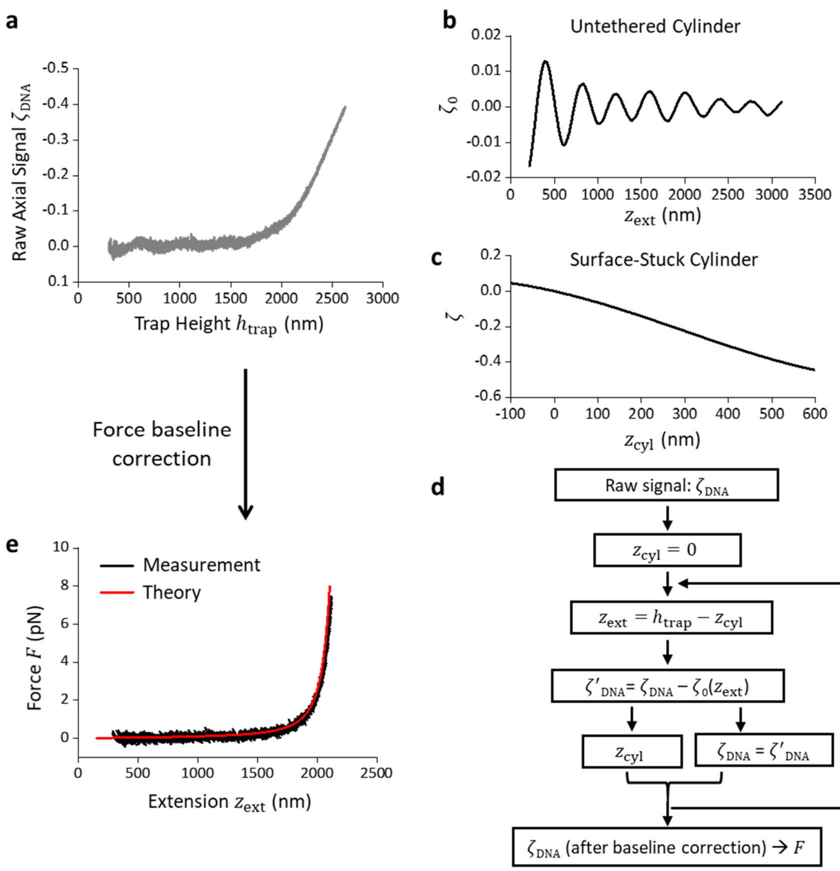


**Fig. S5.** An iterative algorithm for AOT force-offset correction for AOT measurements using the metamaterial elliptical cylinders.

The force to stretch a DNA molecule is measured via the normalized axial displacement detector signal $\zeta_{\mathrm{DNA}}$ with an increase in the trap height $h_{\mathrm{trap}}$: $F = k_z \cdot (-\zeta_{\mathrm{DNA}})/S_z$, where $k_z$ is the trap stiffness, and $S_z$ is the axial displacement sensitivity(*4*). However, this raw signal contains a baseline due to the Fabry-Pérot effect that results from interference between the cylinder bottom surface and the sample chamber surface. The amplitude of the periodic force signal is negligible for the conventional quartz cylinder, whose refractive indices ($n_{\mathrm{e}}$ = 1.54 and $n_{\mathrm{o}}$ = 1.53) are comparable to the surrounding medium ($n$ = 1.326). However, the high refractive indices of the metamaterial elliptical cylinder ($n_{\mathrm{e}}$ = 1.66 and $n_{\mathrm{o}}$ = 1.75) result in a much stronger interference and larger amplitude of the periodic force baseline (**a**).

To correct for this baseline, we performed the following procedure. We first measured the signal baseline $\zeta_0$ of untether metamaterial elliptical cylinders to determine its dependence on the distance between the cylinder bottom surface and the sample chamber surface, which will corresponds to $z_{\text{ext}}$ in a DNA stretching experiment (**b**). We next measured the signal $\zeta$ versus the cylinder displacement $z_{\text{cyl}}$ by axially scanning surface-immobilized cylinders through the trap (**c**). Then, we used an iterative algorithm to convert the raw DNA-stretching data $\zeta_{\text{DNA}}(h_{\text{trap}})$ (**a**) to the corrected force versus extension data (**d**). The resulting force-extension curve agrees well with that prediction, validating this method (**e**). This method was implemented in the real-time DNA torsional measurements under a constant force (Fig. 5; Fig. S6).

**a**. Raw DNA-stretching data $\zeta_{\text{DNA}}$ versus $h_{\text{trap}}$ using a metamaterial elliptical cylinder.

**b**. Signal baseline $\zeta_0$ of untether metamaterial elliptical cylinders and its dependence on the distance between the cylinder bottom surface and the sample chamber surface, mimicking the end-to-end extension $z_{\text{ext}}$ when being attached with a DNA molecule. Shown is an averaged baseline measured with *N* = 6 cylinders.

**c**. Signal $\zeta$ versus the cylinder displacement $z_{\text{cyl}}$ by axially scanning surface-immobilized cylinders through the trap. Shown is an averaged relation measured with *N* = 13 cylinders.

**d**. Iterative algorithm implemented on the AOT for real-time signal correction to achieve precise force and extension measurements.

**e**. DNA force-extension curve after the force-offset correction, which agrees with the prediction using the modified Marko Siggia worm-like chain model(*5, 6*).

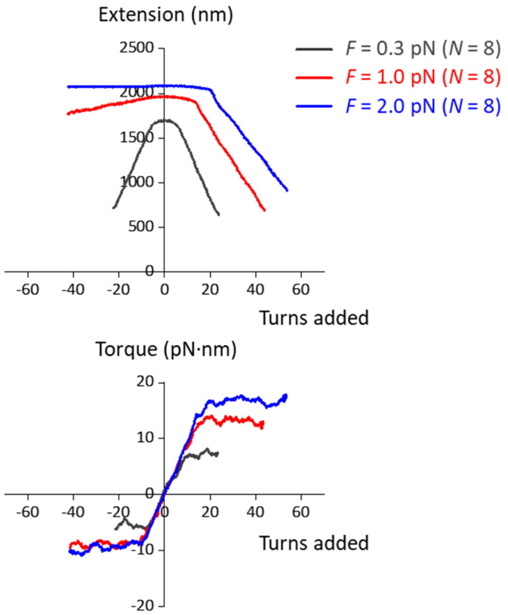


**Fig. S6.** DNA torsional measurements by metamaterial elliptical cylinders under different forces. These measurements show that metamaterial elliptical cylinder can accurately measure the torsional properties of DNA over the standard force range for single-molecule experiments(*2*). The traces shown were averaged from individual measurements of *N* = 8 DNA molecules, where a sliding window of 0.1 turn was applied to average extension and a sliding window of 4 turns was applied to average torque.
